# Compressed phenotypic screens for complex multicellular models and high-content assays

**DOI:** 10.1101/2023.01.23.525189

**Authors:** Benjamin E. Mead, Conner Kummerlowe, Nuo Liu, Walaa E. Kattan, Thomas Cheng, Jaime H. Cheah, Christian K. Soule, Josh Peters, Kristen E. Lowder, Paul C. Blainey, William C. Hahn, Brian Cleary, Bryan Bryson, Peter S. Winter, Srivatsan Raghavan, Alex K. Shalek

**Author notes:** These co-first authors contributed equally (Benjamin E. Mead, Conner Kummerlowe). These co-second authors contributed equally (Nuo Liu, Walaa E. Kattan). These co-senior authors contributed equally (Peter S. Winter, Srivatsan Raghavan, Alex K. Shalek).

## Abstract

High-throughput phenotypic screens leveraging biochemical perturbations, high-content readouts, and complex multicellular models could advance therapeutic discovery yet remain constrained by limitations of scale. To address this, we establish a method for compressing screens by pooling perturbations followed by computational deconvolution. Conducting controlled benchmarks with a highly bioactive small molecule library and a high-content imaging readout, we demonstrate increased efficiency for compressed experimental designs compared to conventional approaches. To prove generalizability, we apply compressed screening to examine transcriptional responses of patient-derived pancreatic cancer organoids to a library of tumor-microenvironment (TME)-nominated recombinant protein ligands. Using single-cell RNA-seq as a readout, we uncover reproducible phenotypic shifts induced by ligands that correlate with clinical features in larger datasets and are distinct from reference signatures available in public databases. In sum, our approach enables phenotypic screens that interrogate complex multicellular models with rich phenotypic readouts to advance translatable drug discovery as well as basic biology.

## Introduction

Phenotypic drug discovery is a powerful approach for identifying clinically relevant treatments^1–5^. To date, successful efforts have employed basic readouts (e.g. viability) in simple models of infectious or monogenic diseases where the underlying mechanisms are easy to faithfully recapitulate^4^. However, most human diseases arise in the setting of complex multigenic and multicellular ensembles that are not well-captured by these approaches. While recent advances in phenotypic screening can now directly link genetic perturbation libraries to high-content readouts (e.g., Perturb-seq^6^), they still have several drawbacks. For example, genetic approaches are highly challenging to use with complex multicellular models and do not mirror the pleotropic manner in which intercellular signals act *in vivo*. Most critically though, any hit cannot be easily advanced as a therapeutic substrate since it provides a gene target rather than a therapeutic agent, disrupting the “chain of translatability”^7^. As a result, there remains a wide array of human maladies for which phenotypic drug discovery cannot be applied effectively. To advance drug discovery efforts, we require improved phenotypic screens that can leverage: 1) *in vitro* or *ex vivo* cellular models that maintain high-fidelity to *in vivo* disease contexts (e.g., cell type, epigenetic state, or tissue of residency); 2) high-content readouts, such as single-cell genomics or high-content imaging, that can comprehensively capture the complex cellular phenotypes associated with a disease state; and, 3) biochemical perturbation libraries built with small molecules or protein ligands that, unlike genetic perturbations, can be directly advanced as therapeutic substrates.

At present, it remains challenging to conduct high-content phenotypic biochemical perturbation screens in complex, high-fidelity avatars due to two limitations of scale. First, high-content readouts, such as single-cell transcriptomics (scRNA-seq), are orders of magnitude more expensive than simple functional assays like cell viability, growth, or secretion. While methods exist for increasing efficiency (e.g., compressed sensing^8^ and antibody or lipid based sample multiplexing^9^), they can still be costly to implement on a per-compound basis and/or have fundamental throughput limitations. Second, high-fidelity models derived from clinical specimens are more challenging to generate at scale than less physiologically representative systems like cell lines. In the most representative cases (e.g., tissue explants or specific model organisms), bio-mass limitations practically restrict the number of perturbations that can be tested; in expandable models of intermediate complexity (e.g., organoids), expansion can be constrained by phenotypic changes over time, as evidenced by epigenetic drift in intestinal organoids^10^, clonal drift in glioblastoma organoids^11^, and altered RNA and genetic state in pancreatic cancer organoids^12^. As one approach to enhance scale, past studies have pooled perturbations together^13^. While this fundamental approach could improve scale and reduce assay costs per-compound, previous efforts have been limited to simple readouts and libraries of mostly inactive small molecules, leaving unanswered questions about the efficacy of such approaches and how to effectively implement them^13^.

Here, we introduce a generalizable, scalable method for compressing phenotypic screens with complex multicellular models and high-content readouts. Our method: 1) pools perturbations to reduce sample number requirements and 2) infers single perturbation effects with a regression-based framework. To validate our method and examine the bounds of compression, we conducted a series of phenotypic screening experiments, both compressed and conventional (ground truth), using a bioactive small molecule library and a high-content imaging readout (Cell Painting)^14^. Across a wide range of pool sizes, we consistently identify the compounds with the largest ground truth effects as hits in our compressed screens, thoroughly vetting the robustness of the approach. We also demonstrate the generalizability of the method to other models and readouts by performing a compressed screen of tumor-microenvironment (TME)-nominated recombinant protein ligands on patient-derived pancreatic ductal adenocarcinoma (PDAC) organoids with a scRNA-seq readout. We find that almost all of our top hits drive conserved transcriptional responses when screened individually. Moreover, we demonstrate the biological importance of these findings by showing that the axes of transcriptional variation we uncover better correlate with important clinical features in larger PDAC datasets whereas current signatures available in reference databases (e.g., MsigDB) do not.

In summary, our method provides a means to conduct high-content phenotypic screening in complex, representative biological models, expanding our ability to connect experimental perturbations with clinical observation to strengthen the “chain of translatability” in drug discovery.

## Results

### Compressed screening: A scalable phenotypic screening framework for complex model systems and readouts

We developed compressed screening to increase the throughput of phenotypic screens by pooling together perturbations. More specifically, to increase scalability, we combine **N** perturbations into pools of unique perturbations of size **P** where each perturbation is repeated in **R** distinct pools (**Fig. 1a-b**). Relative to a conventional screen where each replicate of each perturbation is screened individually, compressed screening reduces the sample number requirements by a factor of **P**, which we refer to as **P-**fold compression. To analyze the output of compressed screens, we developed an assay-independent computational framework for deconvoluting the effects of compounds based on regularized linear regression and permutation testing, inspired by previous work inferring the effects of guide RNAs on genes in pooled CRISPR screens (**Fig. 1c**)^6^. By developing compressed screens, we aimed to unlock the use of higher complexity model systems and assays in phenotypic screening (**Fig. 1d**).

**Figure 1:**
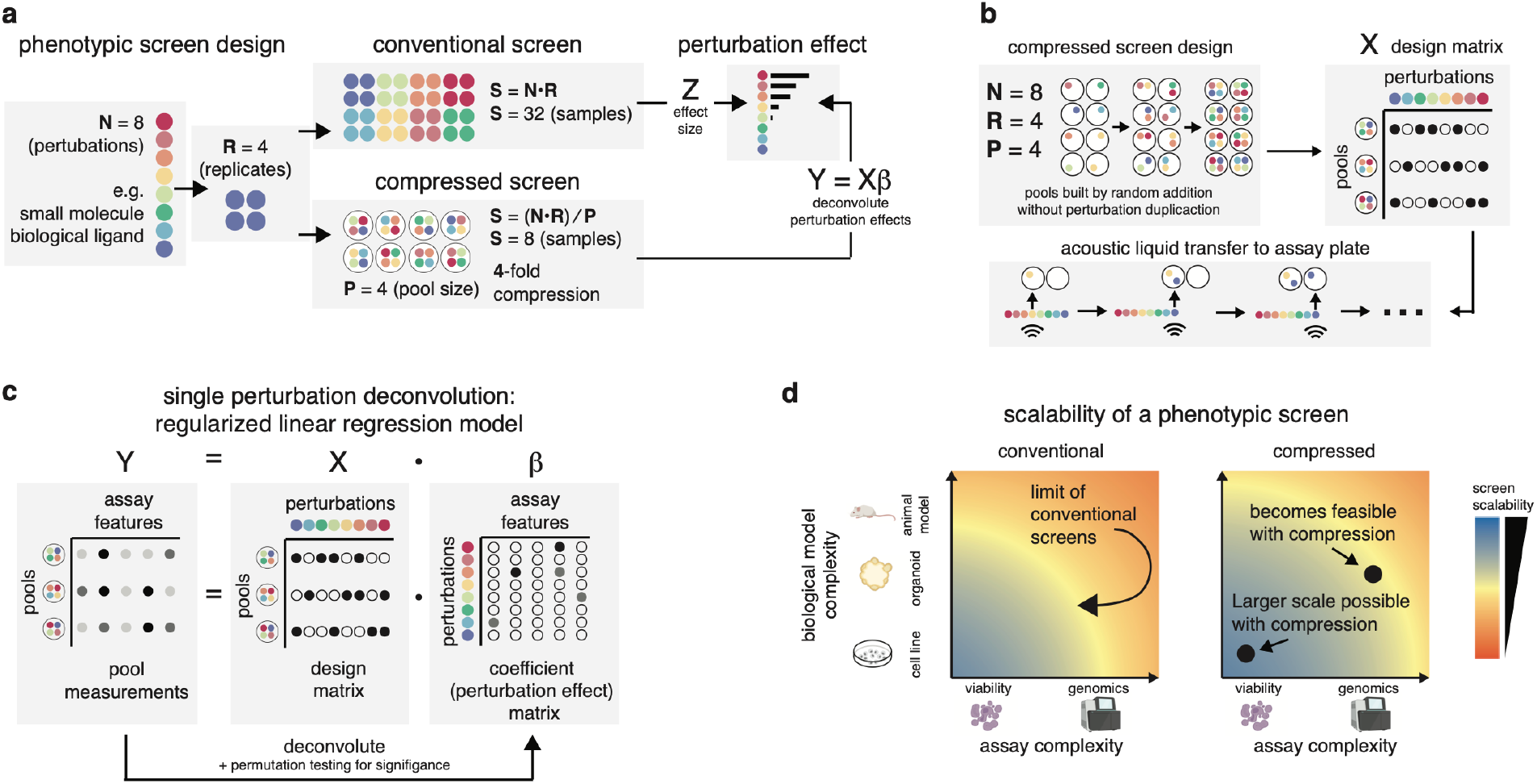
Compressed screening with high-fidelity model systems and high-content assays. **a**, Comparison of the number of samples required to conduct a phenotypic screen in a conventional and compressed manner with N=8 perturbations and R=4 replicates of each perturbation. **b**, Visualization of the construction of a compressed screen with an acoustic liquid handler. **c**, Regression framework for inferring the effects of individual perturbations in a compressed screen: We solve for the coefficient matrix (**β**) that describes the effect of perturbations (whose assignment to pools is denoted in the design matrix **X**) on the measured features of the screen (matrix **Y)**. **d**, Conceptual visualization of how assay and biological model complexity may limit the scalability of conventional screens, as well as how this scalability boundary may be increased in a compressed screen.

### Technology development in U2OS cells lines with a cell-painting readout

To rigorously compare compressed screening with conventional approaches, we conducted a series of ground truth and compressed screens in the U2OS cell line with 316 small molecules from an FDA drug repurposing library using a Cell Painting readout (**Fig. 2a, Table S1**). We selected U2OS, a non-sample limited model system, and Cell Painting, a cost-effective high-content readout, so that we could test the limits of our method by conducting many screens at varying compression levels. We chose an FDA drug repurposing library to test our ability to deconvolute a pooled experiment where highly bioactive perturbations with large effect sizes would frequently co-occur.

**Figure 2:**
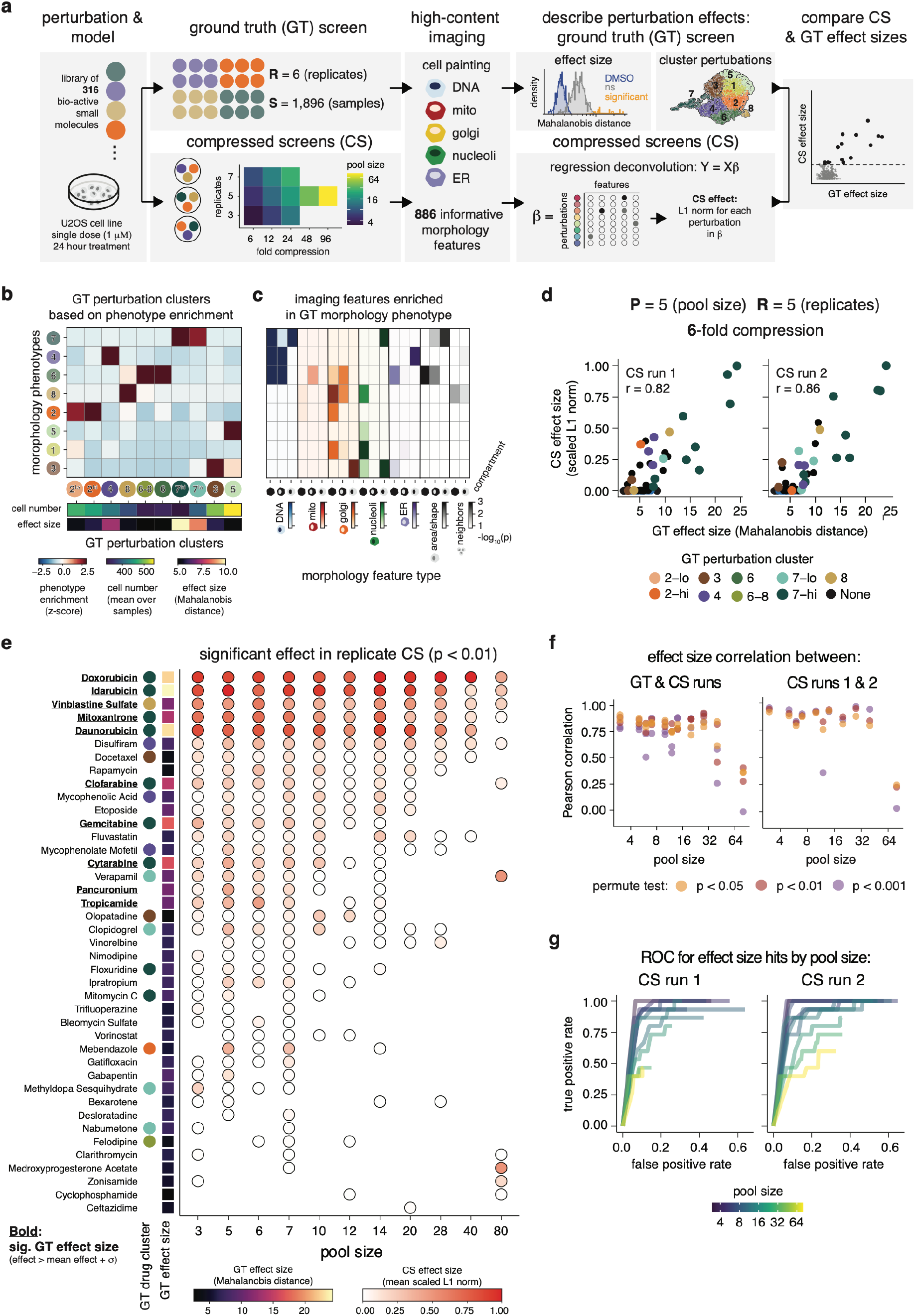
Compressed screening identifies compounds with largest effects in a ground truth setting. **a**, Overview of screens (ground truth (GT) and compressed screens (CS)) and analytical approach for validating the technology and assessing the maximum compression factor that is feasible. **b**, Heatmaps of the GT cellular phenotypes that each GT perturbation cluster is enriched in (fingerprint z-score), as well as the average number of cells per well and Mahalanobis distance for each GT perturbation cluster. **c**, Heatmap of the Fisher’s exact enrichments (-log10(p value)) of the features differentially utilized by each GT phenotype (log2 fold change > 3) in the 7 types of Cell Painting features. Bottom bar visualizes the mean number of cells per well across all samples in each GT phenotype. **d**, Scatterplots of the inferred perturbation effects in a compressed screen (Scaled L1 norm) vs. the GT effect (Mahalanobis distance) for two replicate runs (6X compression, 5 replicates of each perturbation) with distinct pool randomization. r, Pearson correlation, CS run1: p value < 2.2*10^−16^, CS run 2: p value < 2.2*10^−16^). **e**, Dotplot of the mean scaled L1 norm of the perturbations called as hits (scaled L1 norm > 0) in both replicate compressed screens at each pool size, as well as the GT perturbation cluster and GT Mahalanobis distance of each perturbation. **f**, Scatterplot over all pool sizes of the fraction of perturbation hits in the CS screen that were significantly enriched in a biological phenotype in the GT screen, for three permute test significance levels (blue – p value < 0.05, green – p value < 0.01, red – p value < 0.001). **g**, ROC curves for each pool size in both CS screens displaying the changes in the true positive and false positive rates for identifying GT significant perturbations as hits in CS screens that occur when varying the permutation testing threshold in deconvolution from 0 to 1 by steps of 0.01.

First, to find optimal conditions for our compressed screen, we conventionally screened 9 concentration/timepoint combinations. To analyze our Cell Painting dataset, we constructed an analysis pipeline to perform illumination correction, quality control, cell segmentation, morphological feature extraction, and highly variable feature selection (see **Methods**). For each sample well treated with a perturbation or a DMSO negative control, we obtained a vector of informative features that measure cell morphology. With this dataset, we quantified the magnitude of the effects of the perturbations relative to the negative controls by calculating the Mahalanobis distance (i.e., the multidimensional z-score) between each perturbation and the DMSO controls. From the 9 conventional screens, we chose the 24 hour timepoint and 1 mM concentration combination as our reference GT screen, and conducted subsequent compressed screens with these conditions, as these conditions yielded the largest variation in perturbation effect size relative to DMSO control (**Extended Data Fig. 1a-b**).

In addition to identifying the magnitude of the GT drug effects (Mahalanobis distance), we characterized the phenotypes associated with drugs via an unbiased clustering approach. To do so, we first identified the cellular morphological phenotypes in our GT screen by clustering over morphological profiles **(Fig. 2b**). Then, for each drug and each morphological phenotype, we calculated an enrichment score that quantified how enriched the samples treated with a given drug were for a given morphological phenotypes **(Extended Data Fig. 1d)**. Finally, we clustered drugs based on their enrichment scores define GT drug clusters (**Fig. 2c, Extended Data Fig. 1e**)^15^. This revealed 10 clusters of drugs that were distinctly enriched for the underlying cellular morphologies present in the data.

We next conducted an array of compressed screens with pool sizes varying from 3 to 80 perturbations in order to benchmark and test the limits of compression with a highly bio-active perturbation library (**Fig. 2a**). First, we benchmarked our approach for deconvoluting individual perturbations in a compressed screen. For each pooled perturbation, we summarized its effect by calculating the L1 norm (sum of the absolute values) of the permute test significant regression coefficients over all features and scaling by the max L1 norm value we measured. These scaled L1 norm values strongly correlated with the GT perturbation effects for pool sizes of up to 40 perturbations and were also strongly correlated between screens with the same perturbations and pool size but distinct random pooling (**Fig. 2d**). Across all pool sizes our approach identified a subset of significant perturbations (scaled L1 norm > 0) corresponding to the compounds with the largest GT effects, while lower compression (smaller pools) recovered lower effect perturbations (**Fig. 2e**). At lower compressions perturbations from 8 of the 10 GT drug clusters were identified as hits, and even at pool sizes as high of 40, perturbations from 4 of the 10 GT drug clusters were identified as hits. Thus, even at high compression values, our compressed screens identified drugs that drove multiple phenotypes and were not limited to identifying the effect of a single dominant phenotype. By varying the permutation testing significance threshold in our approach, we could tune deconvolution to either identify more hits that drive significant GT phenotypes with a higher false positive rate or to uncover fewer hits but with a lower false positive rate, with pool sizes up to 40 perturbations having strong agreement to GT effect (**Fig. 2f-g**). In summary, our benchmarking results demonstrate that our approach to screening is efficient, accurate, reproducible, and tunable.

### Compressed tumor microenvironment ligand screening with PDAC organoids and scRNA-seq

To evaluate the utility of our approach for additional perturbations, model systems, and readouts, we applied compressed screening to characterize biological ligand responses in patient-derived pancreatic ductal adenocarcinoma (PDAC) organoids using single-cell RNA-seq (scRNA-seq). PDAC tumor cells *in vivo* fall along the prognostically and therapeutically relevant basal-to-classical transcriptional state continuum^16^. scRNA-seq studies have demonstrated that tumors enriched for specific cancer cell states are associated with specific non-cancer cell populations that express genes encoding for biological ligands^17^. However, it is challenging to conduct conventional function screens of these ligand proteins due to the costs of organoid culture and scRNA-seq. Thus, this problem posed an ideal application for compressed screening.

We applied compressed screening to perturb PDAC organoids with a library of 68 recombinant protein ligands nominated by the expression of their corresponding genes in immune and structural cells that are likely to be present in the PDAC tumor microenvironment (TME)^17,18^, and used a multiplexed scRNA-seq readout to measure phenotypic ligands effects (**Fig. 3a, Table S2**)^19^. To test replicability, we conducted two runs with the same model, library, and compression scheme (replicates = 5, mean pool size = 4.75) but distinct randomly generated ligand pools. In addition to negative controls (wells containing only organoids and minimal media), we individually screened three ligands (TNF-α, TGF-β1, IFN-γ) with known activity in PDAC^12,20^ as positive controls (landmarks). After sample demultiplexing and quality control, we analyzed 5,662 cells and 10,881 cells from the two runs, with a mean ± standard deviation of cells per compressed pool of 59 ± 32 cells and 113 ± 65 respectively (**Extended Data Fig. 2a-b**).

**Figure 3:**
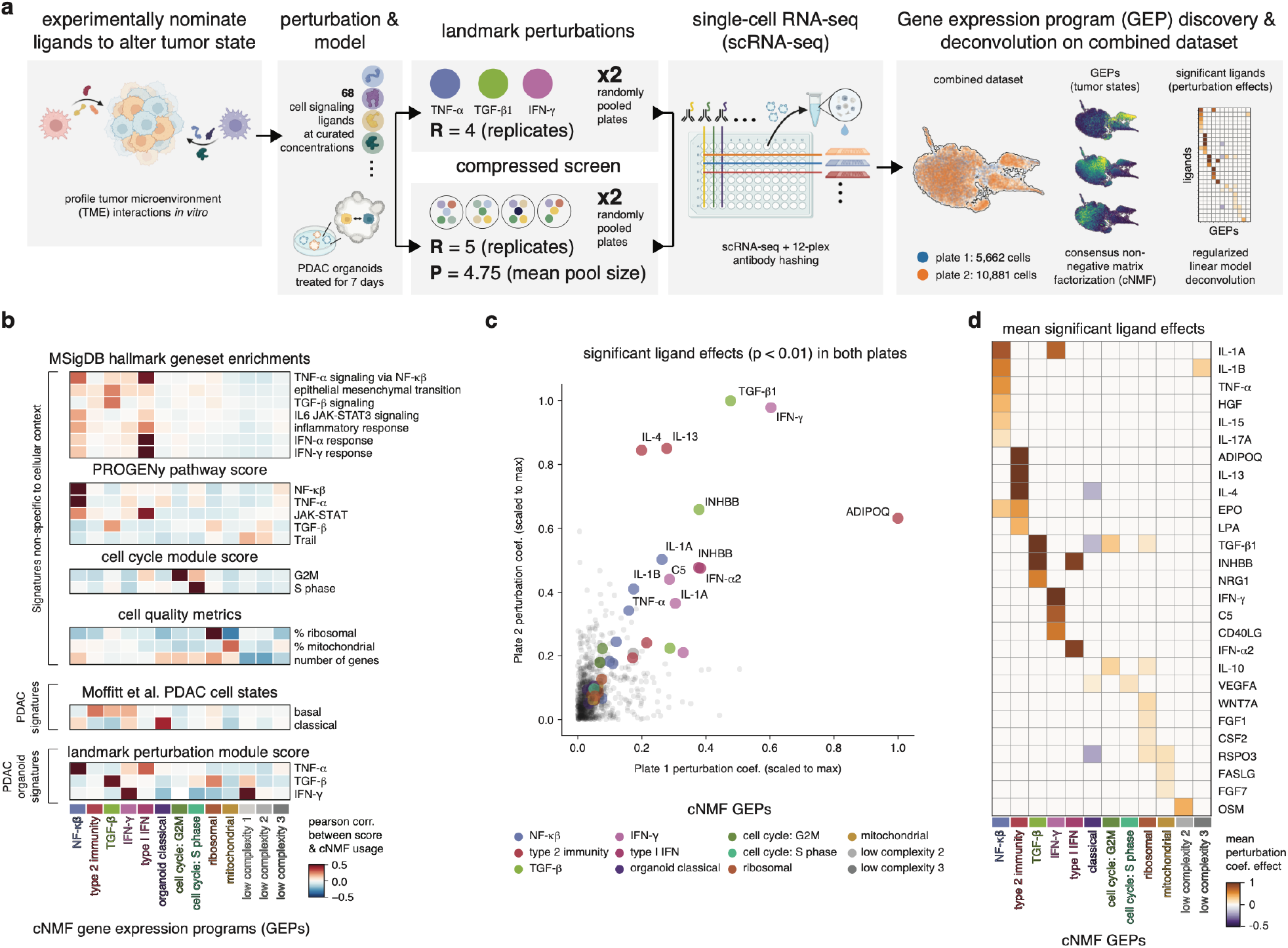
Compressed screen of biological ligands in PDAC organoids reveals major axes of transcriptional response. **a**, Overview of biological ligand compressed screen with PDAC organoids and scRNA-seq analysis approach **b**, Heatmaps visualizing the Pearson correlation across cells of the usage of the cNMF gene expression programs and the module score for existing gene signatures. **c**, Scatterplot of significant ligand – cNMF module effects (deconvolution regression coefficients) from two compressed screens with distinct random pooling. **d**, Heatmap of the mean ligand – cNMF module effect over both compressed screens.

As PDAC cells in these screens were simultaneously exposed to multiple perturbations, it seemed likely that a given cell could concurrently express multiple gene expression programs in response to stimulation. Thus, we applied consensus non-negative matrix factorization (cNMF) to infer the predominate gene expression programs (GEPs) in our data as well as the activity of each GEP in each cell^21^. This revealed 13 GEPs with highly variable activity across cells (**Extended Data Fig. 2c, Table S3**). We annotated 11 of these GEPs by correlating their expression with module scores of existing gene sets from pancreatic cancer cell states, the MsigDB and PROGENy databases, cell cycle scores, and cell quality metrics (**Fig. 3b, Extended Data Fig. 2d-e**)^22^. These 11 gene sets were as follows: NF-κβ activation, TGF-β response, Type I IFN response, Moffitt et al. Classical, cell cycle S phase, cell cycle G2M, mitochondrial, ribosomal, and three GEPs primarily expressed in low complexity cells. One of the remaining GEPs displayed negligible association (R < 0.03) with the Hallmark IFN-γ response in MsigDB, yet clearly correlated with the module score for genes differentially expressed in the IFN-γ positive control wells. The final GEP did not clearly correlate (R > 0.25) with any signature in MsigDB or PROGENY, and we annotated it as a “type 2 immunity” signature based on examining the deconvolution results presented below and discovering that the top ranked genes in the GEP can be induced by IL-4 and IL-13^23–25^.

We next applied our deconvolution framework to infer which ligands drove the activity of each cNMF GEP in the compressed run (**Fig. 3c-d, Extended Data Fig. 2f**). As stated above, we annotated the type 2 immunity GEP based on its association with IL-4 and IL-13, as well as ADIPOQ, in both compressed screens. Reassuringly, we found that IFN-γ was associated with the IFN-γ response GEP, IFN-α2 was associated with the type I IFN signaling GEP, TGF-β and INHBB (a member of the TGF-β superfamily) with the TGF-β response GEP, IL-1A, IL-1B, and TNF-α (known activators of NF-κβ^26^) with the NF-κβ activation GEP, and the mitogens WNT7A and RSPO3 were associated with the ribosomal GEP^27,28^.

Interestingly, we would not have been able to infer which ligands were most active in the organoids by examining cognate receptor expression, given our active ligands spanned the range of cognate receptor expression in negative control organoid cells (**Extended Data Fig. 2g**). Together, these findings reveal major axes of phenotypic variation in PDAC organoids and nominate TME ligands that drive this variation.

### Validation of compressed hits with single-ligand perturbations

To validate the results of our compressed screen, we individually tested the eleven ligands associated with the 5 aforementioned GEPs (NF-κβ, type 2 immunity, TGF-β, IFN-γ, and ribosomal). This subset of ligands consisted of IL-1A, IL-1B, TNF-α, IL-4, IL-13, ADIPOQ, TGF-β1, INHBB, IFN-γ, WNT7A, and RSPO3. We conducted two such validation screens on separate days (biological replicates), screening six replicates of all ligands, including a set of negative control organoids in each run. We then pooled these data with the negative controls and individual landmark ligand positive controls from the compressed screens to form a final single-ligand perturbation dataset (**Fig. 4a, Extended Data Fig. 3a**).

**Figure 4:**
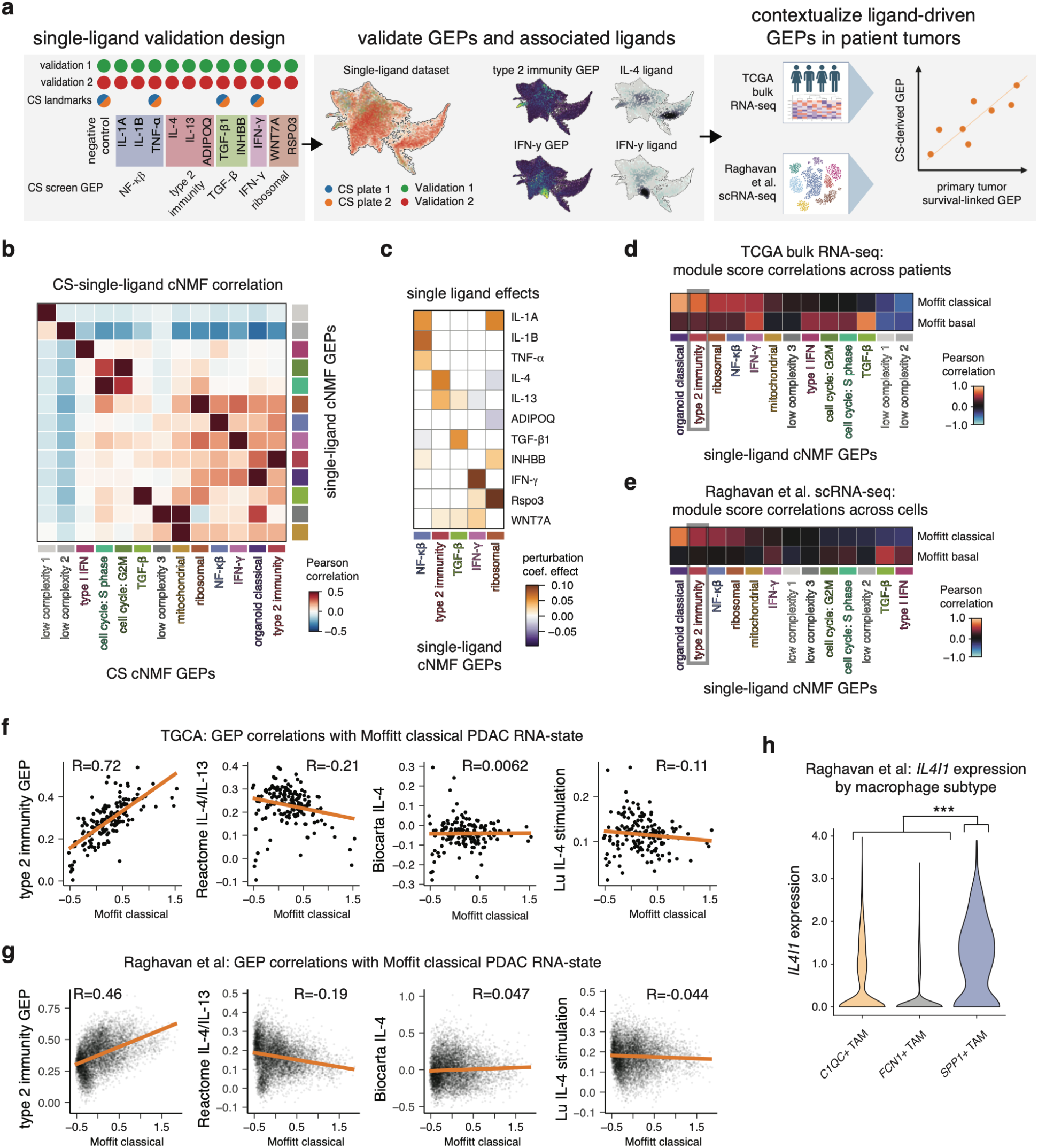
Context specific signatures from compressed screening validate and recontextualize existing primary tumor data. **a**, Overview of single-ligand validation experiments and dataset. **b**, Heatmap of the Pearson correlations of select compressed and single-ligand cNMF modules. **c**, Heatmap of the significant (adj. p value < 0.05) non-zero regression coefficients by ligand for five cNMF modules of interest. **d**, Heatmap of the Pearson correlation across PDAC tumors from TCGA bulk RNA-seq data of the expression of the classical or basal transcriptional states with the expression of each cNMF module. **e**, Heatmap of the Pearson correlation across malignant single cells from PDAC tumors from Raghavan et al of the expression of the **f**, Scatterplots of the correlation of the classical score across PDAC tumors from TCGA bulk RNA-seq with the score of the type 2 immunity GEP and two IL-4 transcriptional signatures from MsigDB. **g**, Scatterplots of the correlation of the classical score across malignant cells in PDAC tumors from Raghavan et al with the score of the type 2 immunity GEP and two IL-4 transcriptional signatures from MsigDB. **h**, Violin plot of *IL4I1* expression in macrophage subtypes in the Raghavan et al dataset.

Running cNMF on this dataset, we identified GEPs that highly correlated with each of the GEPs from the compressed screening dataset (**Fig. 4b, Extended Data Fig. 3b-c)**. When analyzing the data, we observed a batch effect between screens conducted on different days (**Extended Data Fig. 3d**). Thus, to identify the effects associated with each single ligand independent of batch, we first ran cNMF over the single-ligand dataset and then ran a linear model to predict the effect of each ligand on each cNMF GEP while including experimental batch as a covariate (**Extended Data Fig. 3e, Table S4**). This revealed that IL-1A, IL-1B, TNF-α, IL-4, IL-13, TGF-β1, IFN-γ, and RSPO3 significantly increased the expression (adj. p value < 0.05) of the GEPs that they were mostly strongly associated with in the compressed screen (**Fig. 4c**), validating 8 of the 11 ligand effects we tested and revealing ligands that modulated the expression of all 5 of the GEPs selected. These findings demonstrate that our compressed screening approach can successfully identify perturbations in complex cancer models that drive a variety of transcriptomic phenotypes.

### Type 2 immune cytokine GEP expression correlates with prognostic cancer cell state in publicly available transcriptomic datasets

To understand the relevance of the phenotypic shifts we identified to therapeutic development, we examined the expression of the top genes from the GEPs we identified in publicly available PDAC tumor transcriptomic datasets. Doing so, we first calculated a module score for each GEP on bulk RNA-seq samples from PDAC tumors in the cancer genome atlas (TCGA)^29^. Additionally, we scored the TCGA samples on the survival-linked classical and basal states scores from Moffitt et al. Correlating GEP scores with basal/classical scores across the samples in TCGA, we found that, in line with past work, the TGF-β GEP score highly correlated with the Moffitt et al. basal score **(Fig. 4d**)^17^. Surprisingly, the type 2 immunity GEP score (driven by IL-4 and IL-13) strongly correlated with the Moffitt et al. classical score (R=0.72, adj. p value < 10^−24^). In contrast, gene signatures of IL-4 taken from MsigDB that were in different cell types displayed much lower correlations with the classical score (Reactome: r=-0.21, p value < 0.03, Biocarta: r =0.0062, p value < 0.94 Lu et al IL-4: R=-0.1, p value < 0.33) (**Fig. 4f**). We also observed minimal overlap between the type 2 immunity GEP and genes from corresponding signatures from MsigDB, suggesting that the type 2 immunity GEP may be context specific to PDAC and more relevant to the *in vivo* setting than existing MsigDB signatures (**Extended Data Fig. 3f, Table S5**).

As these results may have been confounded by the bulk nature of TCGA samples, we next asked if the same associations held true in malignant cells from the publicly available Raghavan et al. PDAC scRNA-seq dataset^12^. Repeating these analyses across the malignant single cells, we again observed a much stronger correlation (R=0.46), p value < 2.2 * 10^−16^) between the type 2 immunity GEP and the Moffitt et al classical score than we did between the MsigDB signatures and the Moffitt et al classical score (Reactome: r=-0.19, p value < 6.6 * 10^−16^, Biocarta: r =3.6 * 10^−5^, p value < 0.94 Lu et al IL-4: R=-0.04, p value < 9.0 * 10^−5^) **(Fig. 4e**,**g)**. In line with this finding, we also found that *SPP1+* macrophages (which were significantly overrepresented in classical tumors in Raghavan et al) highly expressed *IL4I1*, a downstream target gene directly induced by IL-4, further suggesting that IL-4 signaling may play a role in shaping the classical state-associated TME (**Fig. 4h, Table S6**). These findings suggest that modulating type 2 immune cytokine signaling may be a promising avenue for controlling the plasticity of the prognostic and drug relevant classical PDAC state. This in turn provides an example of the insights that may be gained by a next generation of phenotypic screens that, with compression, effectively leverage complex multicellular models and high-content assays.

## Discussion

Scalability presents a major obstacle to phenotypic screening in complex multicellular models with high-content readouts. Here, to address this challenge, we developed compressed screening – an assay and model independent approach to increase phenotypic screen scalability by pooling perturbations and computationally inferring the effects of individual factors. We developed this method for chemical and biological ligand perturbations, which unlike genomic perturbations, recapitulate *in vivo* intercellular signaling dynamics, are readily advanced as therapeutic substrates, and can be easily used to perturb complex multicellular models. We validated this method in U2OS cells with a high-content imaging readout, demonstrating that the top hits across a wide range of compression corresponded to those perturbations with the largest effects when screened individually. Next, we applied this technology to identify the phenotypic effects of TME-associated ligands in primary PDAC organoids. Here, we discovered multiple classes of ligands that drove conserved patterns of response and, importantly, that IL-4 and IL-13 induced a transcriptional state in PDAC organoid models that not only differed from transcriptomic signatures of IL-4 obtained in other contexts but also may be associated with the clinically relevant classical transcriptional state. Our data suggested that cytokines associated with type 2 immunity may be a mechanism to control the plasticity of this prognostic and drug relevant phenotypic state. Combined, our results demonstrate the broad applicability of compressed screening across models and readouts, as well as the value of leveraging compressed screening to interrogate the phenotypic effects of biological perturbations in a high-throughput manner.

When developing the compressed screening technology in U2OS cells, we found that a linear-model based deconvolution approach could accurately identify compounds with the greatest effects in a ground truth screen across a variety of compressions (3-40 small molecules in each pool). This suggests that non-linear interactions do not confound interpretation until pool sizes are above 40 compounds per pool. This is in line with other work that used high-content imaging to investigate all pairwise interactions between bio-active small molecules and found that only 5% of all possible interactions were non-linear^30^. Nonetheless, in the highly bioactive library we tested, the effects we inferred began to deviate from the ground truth at compression levels above 40 perturbations per pool. This may be due to the fact that, as the pool size increases, the number of possible pairwise interactions between perturbations scales quadratically 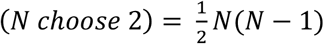, suggesting that even if the fraction of interactions that are non-linear remains small, the absolute number of non-linear interactions will drive a large effect at a large enough pool size. While these findings provide an upper bound for compression with highly bioactive perturbation libraries, much greater compression should be feasible in libraries with fewer bioactive perturbations (e.g., design-of-synthesis (DOS) libraries or discovery compound decks), thereby enabling rapid evaluation of large compound sets in complex biological models. Also, the limit of compression may increase with the richness of the assay readout used in a screen. Additionally, these findings suggest a compelling regime in which to consider searches for combinatorial effects by 1) conducting a compressed screen to find significant individual perturbations and then 2) either screening all pairwise combinations of these significant perturbations or performing a screen in conjunction with one of the factors being held constant to find synergistic pairs.

We further demonstrate that our compressed screening and deconvolution framework is tunable to the needs of the user. By altering the significance threshold for calling a hit in our screens, we show that it is possible to conduct more permissive screens (greater true hit identification along with more false positives) or less permissive screens (fewer true hits but with fewer false positives). This empowers users to adjust their screening strategy based on the scope of planned follow-up validation experiments. While we conducted compressed screens in duplicate with distinct randomization to develop the technology, we observed reproducible top hits between replicates. This suggests that doing a single compressed screen may be sufficient for new lead discovery, albeit with a slightly higher false positive rate than with replicate screens.

Contextualization of signatures identified from our compressed screen in PDAC organoids within clinically relevant datasets suggests that secretion of the type 2 immune cytokines IL4 and IL13 is associated with the PDAC classical transcriptional state. Our prior work demonstrates variation in the PDAC TME that correlates with malignant cell state, with evidence for enrichment of *SPP1+* macrophages in more classical tumors^12^. Notably, *SPP1+* macrophages expressed IL4 target genes, providing additional evidence to suggest increased exposure to IL4 in classical contexts. Taken together, these observations raise the hypothesis that IL4 may be a key state-specific modulator of both malignant cells and their local TME and an important signaling axis for new therapeutic development in PDAC. However, further mechanistic work is needed to dissect the causal relationships between type 2 immune cytokine secretion, classical state expression in malignant PDAC cells, and macrophage recruitment and differentiation within the TME.

In sum, our approach to constructing and deconvoluting compressed screens is assay and model independent and is thus broadly applicable to a variety of model systems and readouts as we have demonstrated here. We anticipate that these compressed screening approaches can be extended to conduct phenotypic screens across a wide array of increasingly complex models (e.g., patient derived organoids, tissue explants, animal models) and readouts (e.g., single-cell genomics, spatial transcriptomics, highly-multiplexed antibody staining (CODEX), multiplexed ion beam imaging (MIBI)) where scalability is a major challenge.

### Limitations of the Study

As designed our technology is intended to serve as a “screening” method and not a “fingerprinting” method. While our technology can identify the perturbations with the largest effects in a library, it cannot perfectly characterize the phenotypic shifts driven by all perturbations in a library. Thus, screens following our approach should be designed with an initial screen to identify top hits, followed by a validation screen that examines the top hits individually. When developing our compressed screens, we only included one concentration for each perturbation; however, in many discovery-oriented applications, the best concentration to use may be unclear. As such, future iterations of this technology may benefit from exploring the feasibility of conducting compressed screens that include multiple doses of the same compound, and deconvolution methods to infer both the effects of, and dose responses to, individual compounds. Furthermore, due to the low frequency at which any two perturbations co-occur, our approach is not optimized to identify non-linear effects within the compressed screen. However, this limitation may not ultimately matter in clinical settings such as chemotherapy, where analysis suggests that the vast majority of combination therapies do not exploit additive or synergistic interactions between compounds, and rather are successful combinations because patient populations are heterogenous and thus the compounds in the combination are effective on distinct subsets of patients or cells within patients^31^.

In summary, compressed screening unlocks complex models and assays which have previously been challenging to leverage for phenotypic screens due to a lack of scalability. This provides new avenues for conducting screens to both map the context specific phenotypic effects of perturbations and develop new therapeutics in systems that are highly relevant to human biology and thus connect *in vitro* discoveries with clinical implementation.

## Acknowledgements

A.K.S. was supported was supported, in part, by the NIH NIDA DP1 Avant-Garde Pioneer Award (1DP1DA053731), the Searle Scholars Program, the Sloan Fellowship in Chemistry, the Bill and Melinda Gates Foundation, the John W. Jarve Seed Fund for Science, and the Ragon Institute. B.B was supported by the NIH (R01A1022553 and R01AR073252). S.R. was supported by Claudia Adams Barr Program for Innovative Basic Cancer Research and the Dana-Farber Cancer Institute Hale Family Center for Pancreatic Cancer Research. W.C.H. and S.R. were supported by the NIH (U01 CA250549).

## Author contributions

Conceptualization, B.E.M., C.K., A.K.S.

Methodology, B.E.M., C.K., N.L., W.E.K. T.C., J.H.C, C.K.S., J.P., B.B., P.S.W. S.R., A.K.S.

Software, B.E.M., C.K., N.L., T.C.

Validation, B.E.M., C.K., W.E.K., P.S.W., S.R.

Formal analysis, B.E.M., C.K., N.L., T.C.

Investigation, B.E.M., C.K., W.E.K., J.H.C., C.K.S.

Resources, J.P., P.C.B., B.B., P.S.W., S.R., A.K.S.

Data curation, B.E.M., C.K., N.L., T.C.

Writing – Original Draft, B.E.M., C.K.

Writing – Review & Editing, B.E.M., C.K., B.C., P.S.W, S.R., A.K.S.

Visualization, B.E.M., C.K.

Supervision, A.K.S.

Project administration, B.E.M., C.K., A.K.S.

Funding acquisition, W.C.H, P.C.B., B.B.. S.R., A.K.S.

## METHODS

### Key resources table

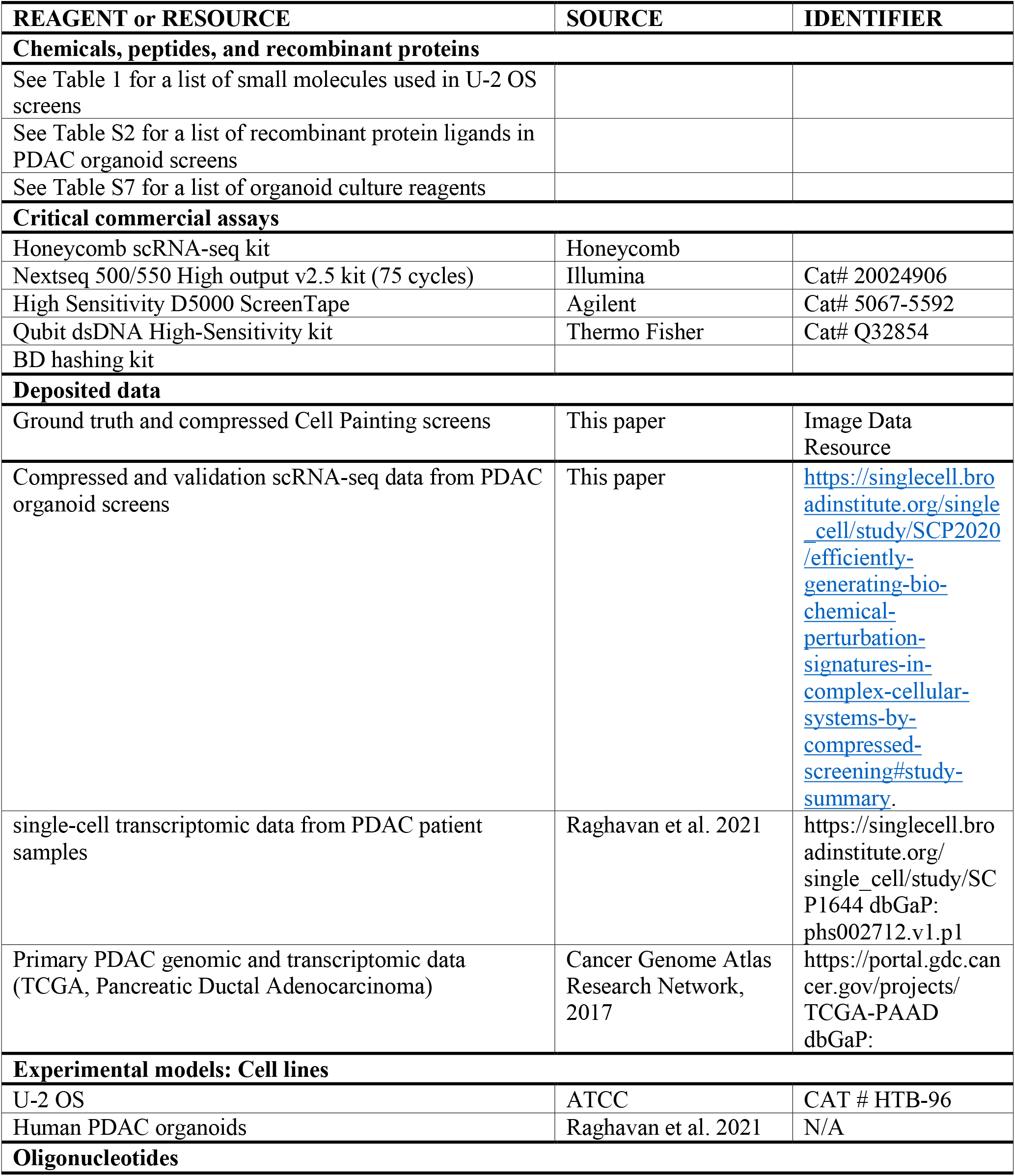
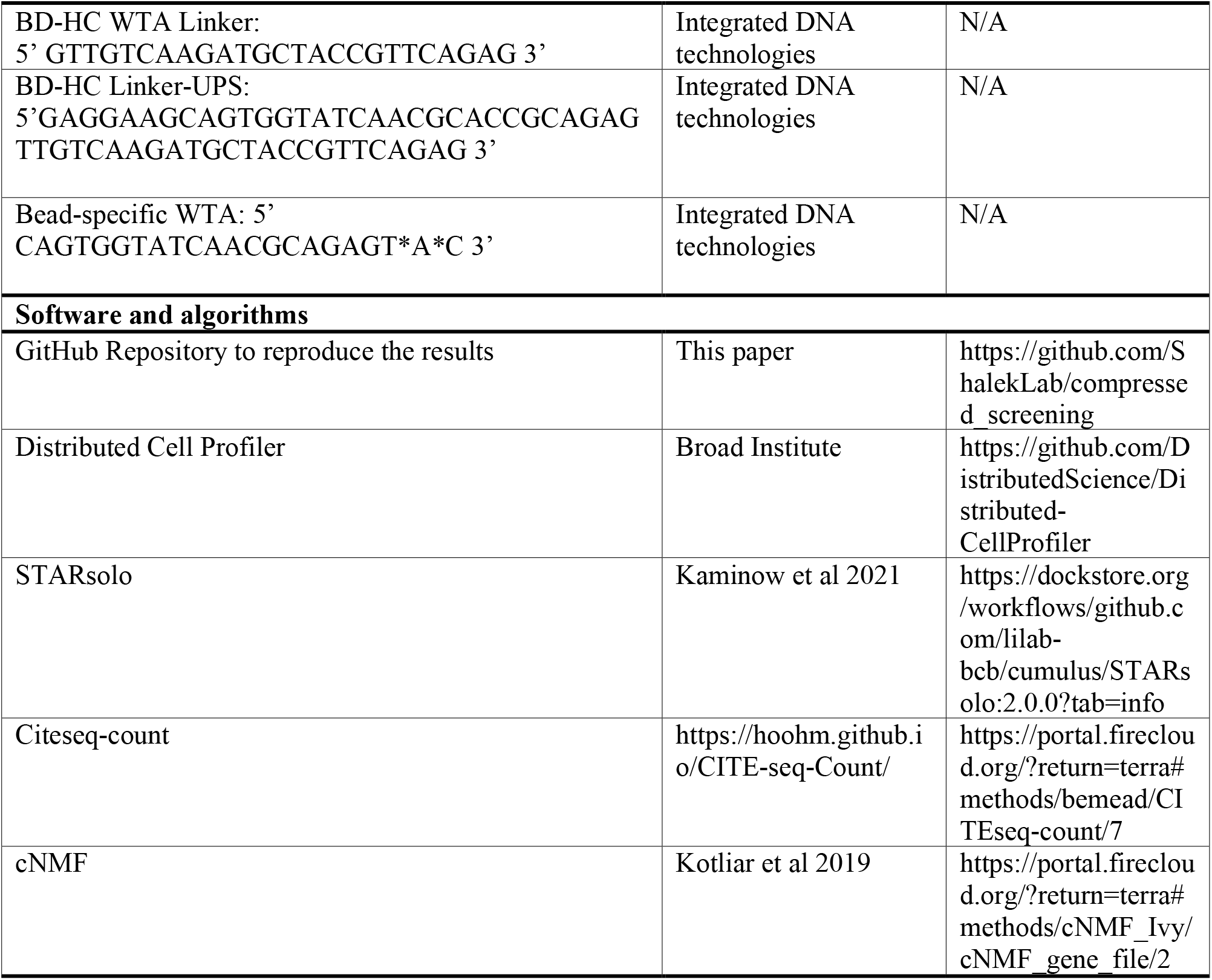

## RESOURCE AVAILABILITY

### Lead contact

Further information and requests for resources and reagents should be directed to and will be fulfilled by the lead contact, Alex K. Shalek.

### Materials availability

This study did not generate new unique reagents. The cell lines, small molecules, and recombinant protein ligands used in this study are available commercially. The organoid model used in this study is available upon request with a materials transfer agreement.

### Data and code availability

- Summarized and deidentified single-cell RNA-sequencing data is available via the Broad Institute’s Single Cell Portal: https://singlecell.broadinstitute.org/single_cell/study/SCP2020/efficiently-generating-bio-chemical-perturbation-signatures-in-complex-cellular-systems-by-compressed-screening#study-summary. Raw sequencing data will be deposited at the NCBI Database of Genotypes and Phenotypes (dbGaP). Links and accession numbers for the publicly available data analyzed in this study are listed in the key resources table.
- Cell Painting images will be deposited in the Image Data Resource.
- Code is available on Github at https://github.com/ShalekLab/compressed_screening.
- Any additional information required to reanalyze the data reported in this paper is available from the lead contact upon request

## EXPERIMENTAL MODEL AND SUBJECT DETAILS

### Cell lines

Culture condition details for U2-OS cells and PDAC organoids are detailed in the methods section describing the screens in which each model was used.

## METHOD DETAILS

### Compressed screen design principles

To design a compressed screen, we utilize a set of user-specified design criteria to randomly assign perturbations to pools while satisfying the following: each pool contains a unique set of perturbations, each perturbation in the library (of size *N* perturbations) occurs in *R*_cs_ replicate pools, and each pool is of average size *P*. The user specifies the library *N*, the compressed replicates *R_cs_*, the conventional replicates *R*_*conv*_, and the desired fold-compression *C*. Fold sample savings of the pools relative to a conventional screen with *S*_*conv*_ samples (*S*_*conv*_ = *N* × *R*_*conv*_ total samples) is *C* compression (*P* = *R_cs_C*/*R*_*conv*_). In each compressed screen, we include a set of negative controls (no perturbation added) and a set of landmark positive controls (treated with individual perturbations known to have large effects). To build the plates for the compressed screen, we transferred perturbations from the source library plates to the pool plates using an Echo 650 Acoustic liquid handler.

### Cell painting experiments

For our cell painting experiments, we plated U20S cells in 96-well black, clear bottom, TC-tread plates (Corning, 3712BC) at 2000 cells per well in 50 µL per well of DMEM (ATCC) + 10% FBS + Pen-Strep media. Cells were then incubated overnight at 37C/5% CO2 to allow cell adhesion and equilibrium. For conventional screening, compounds (316 FDA-approved small molecules, see **Table S1**) were arrayed in 384-well format as stocks that were 1000X the final concentration in DMSO. We then pin transferred (V & P scientific mounted onto an MCA96 head of Tecan Freedom Evo 150) 50 nL of compound into the 50 µL of media per well of the assay plate. Cells were then incubated for 6, 24, or 48 hours at 37C/5% CO2. For compressed screening with the same compound library, we designed screens with the following compression schemes: *C*=6, *R*_*cs*_=3; *C*=6, *R*_*cs*_=5; *C*=6, *R*_*cs*_=7; *C*=12, *R*_*cs*_=3; *C*=12, *R*_*cs*_=5; *C*=12, *R*_*cs*_=7; *C*=24, *R*_*cs*_=3; *C*=24, *R*_*cs*_=5; *C*=24, *R*_*cs*_=7; *C*=48, *R*_*cs*_=5; *C*=96, *R*_*cs*_=5. Respectively, these designs contained 3, 5, 7, 6, 10, 14, 12, 20, 28, 40, and 80 drugs per pool. Furthermore, for each compression scheme, we designed two screens with differing random assignment of drugs to pool. Each plate design was then dispensed via Echo 650 Acoustic liquid handler into 50 µL of media in the 384-well format, where DMSO concentration in negative controls was held constant relative to total compound dispensed for that assay plate. Assay plates were incubated for 24 hours prior to Cell Painting assay.

Following compound treatment, cells were stained for the mitochondria (500 nM MitoTracker Deep Red ThermoFisher #M22425 in DMSO, 7.5 µL stock per assay plate) and for the Golgi apparatus and plasma membrane (60 ug / ml Wheat Germ Agglutinin – AlexaFluor 594, ThermoFisher #W11262 in water, 900 µL stock per assay plate) in 30 µL of media per well and incubated at 37C for 30 minutes. We then fixed the cells by adding 10 µL of 16% methanol-free paraformaldehyde (PFA, ThermoFisher #28908) per well and incubated at room temperature for 20 minutes. Next, we aspirated the solution, washed cells with 70 µL per well of 1X HBSS (Invitrogen, #14065-56), and permeabilized cells with 30 µL per well of 0.1% Triton X-100 in 1X HBSS at room temperature for 20 minutes. We then again washed cells with 70 µL per well of 1X HBSS and stained for the nucleus, endoplasmic reticulum, nucleoli, and F-actin by adding to each plate 1.5 mL of 100 ug/ml ConcanavalinA-AlexaFluor 488 (ThermoFisher, #C11252) in 0.1M sodium bicarbonate, 375 µL Phalloidin-AlexaFluor 594 (ThermoFisher #A12381) in methanol, 7.5 µL of 5 µg/ml Hoeschst33342 (ThermoFisher #H3570) in water, 15 mL of 3 µM SYTO14 (ThermoFisher #S7576), and 15 mL of 1X HBSS in 1% BSA. Cells were then incubated at room temperature for 30 minutes. We then aspirated the solution and washed cells three times with 70 µL of 1X HBSS, and then filled the wells with 70 µL of HBSS, sealed the plate, and stored plates at 4C until ready for imaging. Cells were imaged on an ArrayScan VTI with a 20X objective, using the Hoescht channel to autofocus and set the Z, and autoexposure set for each channel. 9 fields were captured in the middle of each well.

### PDAC organoid culture

Patient-derived pancreatic cancer organoids^12^ were seeded in Growth factor Reduced Matrigel (Corning), fed with human complete organoid medium containing Advanced DMEM/F12 (GIBCO), 10 mM HEPES (GIBCO), 1x GlutaMAX (GIBCO), 500 nM A83-01 (Tocris), 50 ng/mL mEGF (Peprotech), 100 ng/mL mNoggin (Peprotech), 100 ng/mL hFGF10 (Peprotech), 10 nM hGastrin I (Sigma), 1.25 mM N-acetylcysteine (Sigma), 10 mM Nicotinamide (Sigma), 1x B27 supplement (GIBCO), RSPONDIN-1 conditioned media 10% final, WNT3A conditioned media 50% final, 100 U/mL penicillin/streptomycin (GIBCO), and 1x Primocin (Invivogen), and maintained at 37°C in 5% CO2. Media was changed every 6-7 days. Established organoid models were then passaged by dissociation with TrypLE Express (Thermo Fisher) for 30 minutes and reseeded into Matrigel droplets and fresh culture medium.

11 candidate ligands were tested for their effects on cell state: 50ng/mL rHu WNT-7A, 500ng/mL rHu R-spondin 3, 10ng/mL rHu TNF-α, 10ng/mL rHu IL-13, 10ng/mL rHu IL-4, 10ng/mL rHu IFNα, 10ng/mL rHu IL-1A, 10ng/mL rHu IL-1B, 10ng/mL rHu TGF?1, 25ng/mL rHu Adiponectin, and 10ng/mL rHu Activin A (Peprotech). The same batch of organoids used in the original screen (PANFR0562) were used for the validation experiment all of which were cultured and maintained in “OWRNA” media: complete <u>o</u>rganoid medium without <u>W</u>NT3A, <u>R</u>SPONDIN-1, <u>N</u>OGGIN, and <u>A</u>-8301 consisting of Advanced DMEM/F12 (Thermo Fisher), 10 mM HEPES (Thermo Fisher), 1x GlutaMAX (Thermo Fisher), 50 ng/mL mEGF (Peprotech), 100 ng/mL hFGF10 (Peprotech), 10 nM hGastrin I (Sigma), 1.25 mM N-acetylcysteine (Sigma), 10 mM Nicotinamide (Sigma), 1x B27 supplement (Thermo Fisher), 100 U/mL penicillin/streptomycin (Thermo Fisher), and 1x Primocin (Invivogen). Organoids were dissociated to single cells using 1X TrypLE Express (Thermo Fisher) for 30 minutes, counted and seeded in suspension (10% Matrigel, 90% media) at a density of 40,000 viable cells/well in a 96-well plate at a volume of 50uL/well. Ligands were added after 24 hours at 2x the desired final concentration in 50uL to bring the total volume per well to 100uL at 1x ligand concentration. Organoids were collected after 7 days for single-cell RNA sequencing. Treated organoids were tested in duplicate while control organoids (treated with 0.1% BSA in 1X PBS as vehicle control) were assessed in quadruplicate. Candidate ligands and their concentrations can be found in Table S1.

### Ligand library selection

To generate a library of relevant, effective ligands, we compiled multiple ligand-receptor datasets ^18,32,33^. We first prioritized ligands expressed by structural cell types in a variety of organs targeting myeloid cells or ligands expressed by human monocyte-derived macrophages or human macrophages *in vivo*. These 200+ ligands were then filtered based on their overall expression and applicability *in vitro* as a secreted, independent effector molecule. Additional non-genetically encoded molecules were added based on well-established effects, such as LPS-EK, LTB4, ox-LDL, and Pam3CSK4. Manual curation and extensive literature research established the final library and concentrations based on previously reported concentrations and results.

### Ligand screening

To perform compressed ligand screening on PDAC organoids, cells were expanded as described, resuspended in OWRNA media with 10% v/v Matrigel, and seeded at 20,000 single cells in 35 µL / well into a flat bottom ultra-low attachment 96-well assay plate (Corning #3474) one day prior to dispensing ligands. A compressed screen was designed by randomly assigning ligands to 68 ligands (**Table S2**) to 72 pools such that each pool contained 4 or 5 ligands (average of 4.75 ligands per pool). Two such plates were designed, with distinct random assignment of ligands to pools in each plate. Ligands were first dispensed into 25 µL / well of OWRNA media in a 384-well format transfer plate (quadrant-wise). After ligand dispense, each well was backfilled with an additional 50 µL / well of OWRNA media. 65 µL / well was then transferred with a Tecan Freedom Evo 150 from each transfer plate quadrant into the 96-well assay plate with PDAC organoids, making a final total volume of 100 µL / well in the assay plate. Ligand concentration was modified to account for the multi-stage transfer such that final assay plate concentrations are as reported (see Supplemental **Table S2**). Assay plates were cultured for 7 days at 37C/5% CO2 and processed for single-cell RNA-seq with cell hashing as follows.

### PDAC single-cell RNA-seq with cell hashing

PDAC organoids for compressed screens and validation experiments were dissociated to single cells, hashed with antibody-derived oligo tags, pooled and captured for single-cell RNA-seq via Honeycomb Bio HIVEs. Briefly, 200 µL of RP-10 (RPMI – ThermoFisher #12633-012 + 10% FBS - ThermoFisher #26140) was added to each well of PDAC organoids in the 96-well assay plate (300 µL total volume) with vigorous mixing. The full volume was then transferred to the upper chamber of a 30-40 µm 96 well filter plate (Pall #8027) situated on a vacuum manifold (Honeycomb Bio). Media was aspirated under vacuum, with organoids larger than the filter pore size retained in the upper chamber, an additional 300 µL of RP-10 was added, mixed, and aspirated under vacuum to remove residual Matrigel and cell debris. The bottom of the filter plate was blotted dry and sealed with a plate seal (ThermoFisher #AB0558), and the upper chamber was resuspended with 100 µL of pre-warmed TrypLE Express (ThermoFisher #12604013), mixed 10-15 times, and incubated for 20 minutes at 37C with intermittent mixing. Digestion was then quenched with the addition of 200 µL RP-10 + 50 µg/mL DNase (Stem Cell Technologies), and the single cell suspension was collected by centrifugation (300g, 5 minutes) into a 96-well u-bottom ultra low attachment plate (Corning #7007). Quenching solution was aspirated and 50 µL / well of cell staining solution (45uL cell staining buffer - BD #554656 + 5uL human multiplexing antibody – BD #633781) was added to each well and incubated at room temperature for 20 minutes. Human multiplexing antibodies were distributed column-wise with each column having a unique tag. Following staining, cells were washed in triplicate by adding 300 µL cell staining buffer, pelleting by centrifugation (400g 5 minutes), and aspirating wash solution. Cells were then resuspended in a volume of cell staining buffer such that when samples are collapsed row-wise (containing non-overlapping tags) the total sample volume was 200 µL. Each row was then treated as a sample for loading onto a Honeycomb HIVE.

Honeycomb HIVE single-cell RNA-seq was performed as described in their October 2021 protocol (v21.10), with modifications to accommodate cell hashing. Briefly, each cell suspension was counted by hemocytometer and normalized to a concentration of 15,000 cells in 1 mL of RPMI. Cells were loaded into the HIVEs by centrifugation at 30g for 3 minutes, and stored in cryopreservation solution at −20 C until library preparation (no longer than 2 weeks). HIVEs were thawed, sealed, lysed and hybridized following the standard protocol. Bead recovery was followed by first strand synthesis (reverse transcription), bead clean up (exonuclease treatment), and second strand synthesis following the standard protocol. Whole transcriptome amplification was modified with the addition of 2.5 µL of 10 µM BD-HC WTA linker primer per 400 µL reaction preparation, and PCR was run following the standard protocol. Following PCR, 15 µL of each WTA reaction per HIVE was pooled size selected with a 0.65X SPRI bead (Beckman Coulter #A63881) addition. The bead bound fraction (cDNA) was washed with a 0.65X SPRI concentrate (Honeycomb Bio), eluted, and used in index PCR per the standard protocol. The supernatant (cell hashing library) had additional SPRI beads added to bring the final ratio to 1.2X, product was eluted from the beads, and run through a round of PCR (10 cycles of the standard WTA PCR, with 2.5 µL each of 10 µM BD-HC linker-UPS and 10 µM Bead-specific primers per 50uL reaction preparation) to append the Linker-UPS sequence to the cell hashing library. This product was then used as input to the standard index PCR protocol. Following index PCR, cDNA product was cleaned via SPRI beads and quantified per the standard protocol. The cell hashing library was size selected with a reverse 0.65X SPRI selection, followed by a 1.2X SPRI pull down, and a second round of reverse 0.65X SPRI selection and 1.2X SPRI pull down, and quantified per the standard protocol. Quantified libraries were pooled and sequenced on either Illumina NextSeq 500 or NextSeq 2000 with HIVE custom read and custom index primers (Honeycomb Bio) with Read 1: 25 Read 2: 50 Index 1: 8 Index 2:8 bases.

## QUANTIFICATION AND STATISTICAL ANALYSIS

### Deconvolution of compressed screens with regularized linear regression

To deconvolute the effects of individual perturbations in a compressed screen, we adopted a regularized linear regression approach that was previously used to infer the effects of guide RNAs on gene expression in pooled CRISPR screens ^6^. This general framework to deconvolution required a design matrix **X** (pools by perturbations) and an assay readout matrix **Y** (assay readout by pools) which we fit with a linear model with elastic net regression in order to infer the coefficient matrix **ß** describing the association between perturbations and assay features. We fit the regression and used crosss validation to determine the optimal l1_ratio using the MultiTaskElasticNetCV function in sklearn with possible l1_rations varying from 0.01 to 1. We then permuted the perturbations labels 1000 and re-ran elastic net regression in order to obtain a null distribution for the perturbation-feature coefficients. We then filtered out any coefficients in our non-permuted regression that occured more than p_value_threshold*1000 times in the permuted data.

### Cell painting data preprocessing

We used the Distributed Cell Profiler software on AWS to process the Cell Painting images https://github.com/CellProfiler/Distributed-CellProfiler. Our pipeline first applied an illumination correction to each image based on per-channel measurements and applied intensity correction on each channel between batches based on differences in the batch-averaged median channel intensity. We next adopted an existing approach for using the isolation forest outlier detection algorithm implemented in sklearn to remove outlier images confounded by artefacts ^30^ based on the whole-image measurement of 97 features. This was performed batch-wise and excluded 4.6% of ground truth batch 1 and 2, 2.0% of compressed batch 1, 2.2% of compressed batch 2, and 1.7% of compressed batch 3 images. We next segmented individual cells, nuclei, and cytoplasm and measured a total of 3,402 morphological features across all five imaging channels and compartments. Each feature was then median aggregated over all cells within a given well, constituting a single sample. We then removed empty wells and wells with fewer than 50 segmented cells and scaled each feature by plate using the RobustScaler function in sklearn centered on DMSO wells and scaled by all wells within each plate. Finally, we selected robust and variable features within a dataset (either dose-time analysis or our ground truth and compressed screen analysis) by retaining features with low noise and high signal. We accomplished this by first calculating the median absolute deviation (MAD) for each feature within a plate. Low-noise features were all features with a MAD greater than the 10^th^ percentile across greater than 90% of the plates in a dataset – features which consistently showed at least some variability in most plates. Within the dose-time analysis 1,859 features passed the low-noise criteria, while in the ground truth and compressed screen analysis 2,117 features passed. High-signal features were all features with a MAD greater than the 90^th^ percentile across greater than 10% of the plates in a dataset – features which showed the most variability in at least some of the plates. Within the dose-time analysis 1,254 features passed the high-signal criteria, while in the ground truth and compressed screen analysis 1,054 features passed. The final feature set was then taken as the intersection of low-noise and high-signal feature sets, for the dose-time analysis this resulted in 861 final features, and in the ground truth and compressed screen analysis it was 886 final features.

### Identifying dose and timepoint for cell painting screens

To determine the dose and timepoint where the compounds in our library had the most activity we screened all compounds in our library in duplicate at nine dose timepoint pairwise combinations (0.1 µM, 1µM, 10 µM; 6 hours, 24 hours, 48 hours). We then quantified the effect of each perturbation we calculated the Mahalanobis distance *D*_*p*_between the mean vector for each perturbation and the negative control (DMSO) samples at the same dose and timepoint. Mahalanobis distance is similar to a multidimensional generalization of a z-score and quantifies how many standard deviations a vector 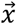 is away from a distribution with mean 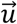 and covariance *S*:

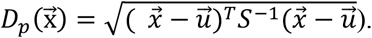

We then calculated the coefficient of variation of the Mahalanobis distance values at each dose/timepoint in order to the dose/timepoint with the broadest range of effects. This occurred at a dose of 1µM and a 24 hour timepoint. We then screened 4 more replicates of each drug at this dose and time point to create a final ground truth dataset consisting of 6 replicates of each drug, which we used to recalculate the Mahalanobis distance between each drug and the negative control distribution. We refer to this dataset as our ground truth (GT).

### Identification of cellular phenotypes in ground truth cell painting data

To identify the biological cellular morphological phenotypes in the GT dataset, we first needed to account for substantial batch effects that existed between the first two replicates in our GT dataset (obtained during the dose-time analysis) and the final four replicates that we collected later. These batch-effects were consistent with large batch effects observed in previous studies using Cell Painting ^34^. To account for this technical variation, we applied Harmony – a batch correction technique that utilizes soft k-means clustering to identify dataset specific batch effects – to our ground truth datasets. After running Harmony, we identified the number of Harmony principle components (PCs) that explained 90% of variance in the data and used these top PCs as the features for identifying the cellular phenotypes in the ground truth data.

To identify cellular phenotypes, we followed the following workflow for clustering the GT cell painting samples. First, we computed the neighborhood graph for all GT samples over the Harmonized features, using the pp.neighbors function in scanpy and a nearest neighbor value of 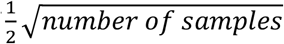. We then clustered on this neighborhood graph using the Leiden algorithm implemented in the tl.leiden function in scanpy. To determine clustering resolution, we re-ran clustering over resolution values from 0.01 to 2.0 and identified the resolution value (0.65) where greater than half (∼52%) of DMSO samples resided in a single cluster, each cluster had significant perturbations, and few perturbations (<0.5%) were significant in more than one cluster (based on phenotypic fingerprinting described below). This identified 8 clusters (GT phenotypes) in the GT cell painting data. For each cluster, we then used the tl.rank_gene_groups function in scanpy to run a Wilcoxon test over the processed and feature selection cell paining features to identify features enriched in each cluster.

To determine perturbations with significant effect we employed an approach for identifying the “phenotypic fingerprints” of the perturbations ^15^. For each perturbation, we counted the number of samples in each of the 8 phenotypic clusters, and then we randomly permuted the perturbation labels 10,000 times and recalculated the samples in each cluster. Using these randomly permuted labels as a null distribution, we calculated the z-scores and p values for the true counts of samples in each cluster. We filtered these results to perturbations with significant enrichment values (maximum z-score > 2 and maximum p value < 0.01). We then clustered the perturbations over these enrichments z-scores and found 10 clusters of perturbations (GT perturbation clusters).

### Compressed cell painting screen design, deconvolution, and comparison to ground truth

To test the limitations of compressed screening, we conducted cell painting screens at a wide array of two parameters: replicates *R* (the number of pools in which and individual drug occurs) and compression *C* (the fold sample savings in the compressed screen relative to our 6 replicate ground truth data). We tested *R* values of 3, 5, and 7, and *C* values of 6, 12, 24, 48, and 96. This resulted in eleven different parameter schemes consisting of 3, 5, 6, 7, 10, 12, 14, 20, 28, 40, and 80 drugs per pool. For each parameter scheme we conducted two screens that were designed with distinct pool randomization which we refer to as CS Run1 and CS Run 2

To deconvolute the compressed screening data, we passed the preprocessed & features selected cell painting features in our general regression framework for deconvolution. To assess the total magnitude of the inferred effect of each perturbation, we then calculated the L1 norm of the regression coefficients across all features for each perturbation. We then calculated the Pearson correlation between these values across all compound and the GT Mahalanobis value in order to assess how well the inferred compressed effects agreed with the effects of perturbations in the ground truth data.

### scRNA-seq alignment, hash demultiplexing, and initial quality control

For each sequencing run, raw sequencing reads were converted from bcl files to FASTQs using bcl2fastq (v2.2) based on Honeycomb Bio indices that corresponded to individual samples. Demultiplexed FASTQs were then aligned to the human GRCh38 genome using STARsolo as implemented by Cumulus (v2.0.0) on Terra.bio. We implemented a 2-pass alignment to first generate a pseudo-whitelist (15,000 cell barcodes with the greatest number of unique molecular identifiers - UMIs) and then realign with this pseudo-whitelist for cell barcode collapse. Additionally, in alignment we specified the following parameters: Multi-mappers: EM, UMI Dedup: 1MM_CR, Solo Features: FullGene, UMI Filtering: MultiGeneUMI_CR, Cell Filter: CellRanger2.2 3000 0.99 10. To align the cell hashing libraries, we provided as input to citeseqcount v1.4.3, implemented via a Terra.bio pipeline, the demultiplexed FASTQs, the pseudo-whitelist from the first pass STARsolo alignment and the hash barcodes per experiment.

Following alignment of sample transcriptomes and cell hashing libraries we performed initial QC filtering, and sample demultiplexing via cell hashing. All samples were filtered to retain only cell barcodes with greater than 500 unique genes and 500 UMIs, and less than 75,000 UMIs and 50% mitochondrial transcripts. Cell barcodes were further filtered by hashing libraries such that only barcodes with greater than 10 hash UMIs, and a 1.1 signal-to-noise ratio (maximum ADT UMI count divided by the second highest ADT UMI count) were retained. Next samples were demultiplexed and labeled by original sample identity by iterating across a range of quantile parameters (0.85 to 1 - (1×10^−6^)) in the Seurat HTODemux function (v4) and choosing the quantile parameter maximizing the number of singlets. Retaining only labeled singlets, dataset were filtered to retain genes detected in at least 10 cells, and used for subsequent analysis.

### PDAC compressed screen scRNA-seq analysis

We next ran consensus non-negative matrix factorization (cNMF) on the combined counts matrices from both compressed runs^21^. We choose an optimal K value, number of cNMF gene expression programs (GEPs), by combining a data driven approach with existing knowledge of PDAC biology. We tested K values ranging from 5 to 50 and then identified candidate optimal K values by plotting the tradeoff between stability and error and focusing on the K values at local stability maxima. We then examined the GEPs at each of these K values and found that at higher K values, a distinct TGFbeta (a known major driver of PDAC biology and one of the landmark positive controls in the screen) GEP was present but at lower values the TGFbeta signaling GEP was collapsed into other GEPs. Thus, chose the smallest of the candidate K values as which we observed a clear TGFbeta signaling GEP.

We proceeded to analyze the combined scRNA-seq dataset in scanpy using the highly variable genes output from cNMF and Pearson residual normalization. We visualized the full dataset by running PCA with the number of PCs chosen by running KneeLocator in python to find the knee of the variance explained by the PCs then running Harmony to integrate the data by batch for visualization purposes only before visualizing the data in an UMAP projection.

We next correlated the usage of the cNMF GEPs across cells and found that a subset described basal expression states that were broadly expressed across all cells were as the remaining were variably expressed across cells. We focused our downstream analysis on these variably expressed cNMF programs. To annotate these GEPs, we first used decoupler and the score_genes function in scanpy to generate module scores for all PROGENy and MsigDB genesets, for the genes differentially upregulated in the TNFa, IFNG, and TGFB1 landmark samples (got this by running scanpy rank genes groups and then running kneelocator), for the top 25 genes in the Basal and Classical signatures from Moffitt et al ^16^, fpr gene signatures for cycling cells and for scRNA-seq quality metrics (nUMI, nFeature, percent of mitochondrial reads, percent of ribosomal reads) ^35^. We then correlated all of these module scores with the GEP usages across all cells and where GEPs highly correlated with a given module, we used this information to annotate the GEPs. With this approach, we annotated all but one highly variable GEP.

Then we passed the usage scores for each highly variable cNMF GEP across cells in the compressed wells into our deconvolution code to infer the effect of perturbations on each GEP. We used these results to annotate the GEP that we could not annotate based on gene signatures.

Ligand receptor analysis: We took the gene names for the 68 ligands in the compressed screen and searched against the CellphoneDB database using the command line call *cellphonedb query find_interactions_by_element*. In siutations were the found interacting partner for a ligand was a complex, the complex was expanded to the individual components.

### PDAC single ligand validation screen scRNA-seq analysis

We adopted the same approach to run cNMF, processes scRNA-seq data, and identify and annotate highly variable GEPs and annotate in the single-ligand data. We then correlated the gene spectra scores in these GEPs to those from the compressed data and focused our analysis on the GEPs that highly correlated with the compressed data. As we observed a large difference in the mean Moffit Classical score in negative control samples from the three different batches of samples in the single-ligand dataset, we assessed the effect of ligands on each GEP by running a linear regression with batch included as a covariate in order to identify ligand effects independent of batch.

### Projecting NMF GEPs onto existing PDAC tumor datasets

To project the validation cNMF GEPs onto existing PDAC tumor datasets from TCGA and Raghavan et al, we first generated a representative genesets for each GEP by running KneeLocator in python on the sorted gene spectra scores to identify the top genes contributing to each GEP. We then generated a module score for each of these genesets, the top 25 Classical and Basal genes from Moffit et al, and IL-4/IL-13 signaling genesets from MsigDB in both PDAC tumor data sets (for each tumor in TCGTA and each cell in Raghavan et al). We then correlated the GEP module scores and MsigDB module scores with the Moffit et al module scores. To look for further evidence of IL-4/IL-13 signaling associated with Classical tumors in Raghavan et al, we did a targeted analysis of the genes differentially expressed in *SPP1*+ macrophages, as this cell population was in significantly higher abundance in the TME of Classical tumors in Raghavan et al.

---

**Extended Data Figure 1:**
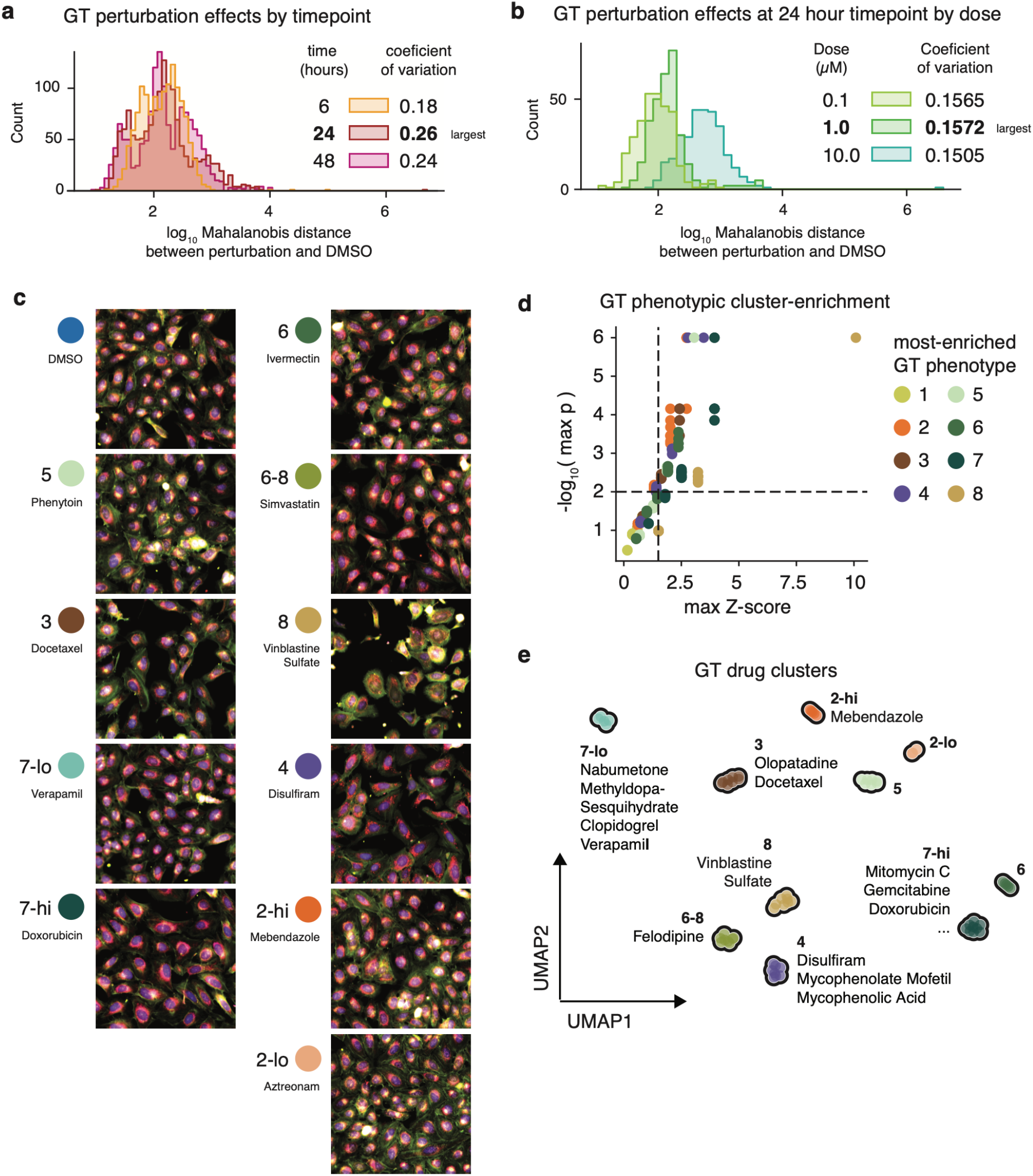
Developing compressed screening by screening 316 small molecules in the U2Os cell line with a Cell Painting readout. **a**, Histogram of the log Mahalanobis distance between each small molecule perturbation and the mean of the distribution of negative control cells (DMSO) at 6 hours, 24 hours, and 28 hours. For each time point, the coefficient of variation of the log Mahalanobis distances (mean / std. deviation) is reported to assess how broad the range of effects is. **b**, Histogram of the log Mahalanobis distance between each small molecule perturbation and the mean of the distribution of negative control cells (DMSO) for the 24 hours timepoint at three doses: 0.1, 1, and 10 µM. For each dose, the coefficient of variation of the log Mahalanobis distances (mean / std. deviation) is reported. **c**, Composite cell painting images from each GT perturbation cluster in the GT screen as well as from top hits from the CS screen. **d**, Scatterplot of non-zero enrichment scores for each perturbation in each GT phenotype **e**, UMAP of all samples in the GT dataset colored by GT perturbation cluster.

**Extended Data Figure 2:**
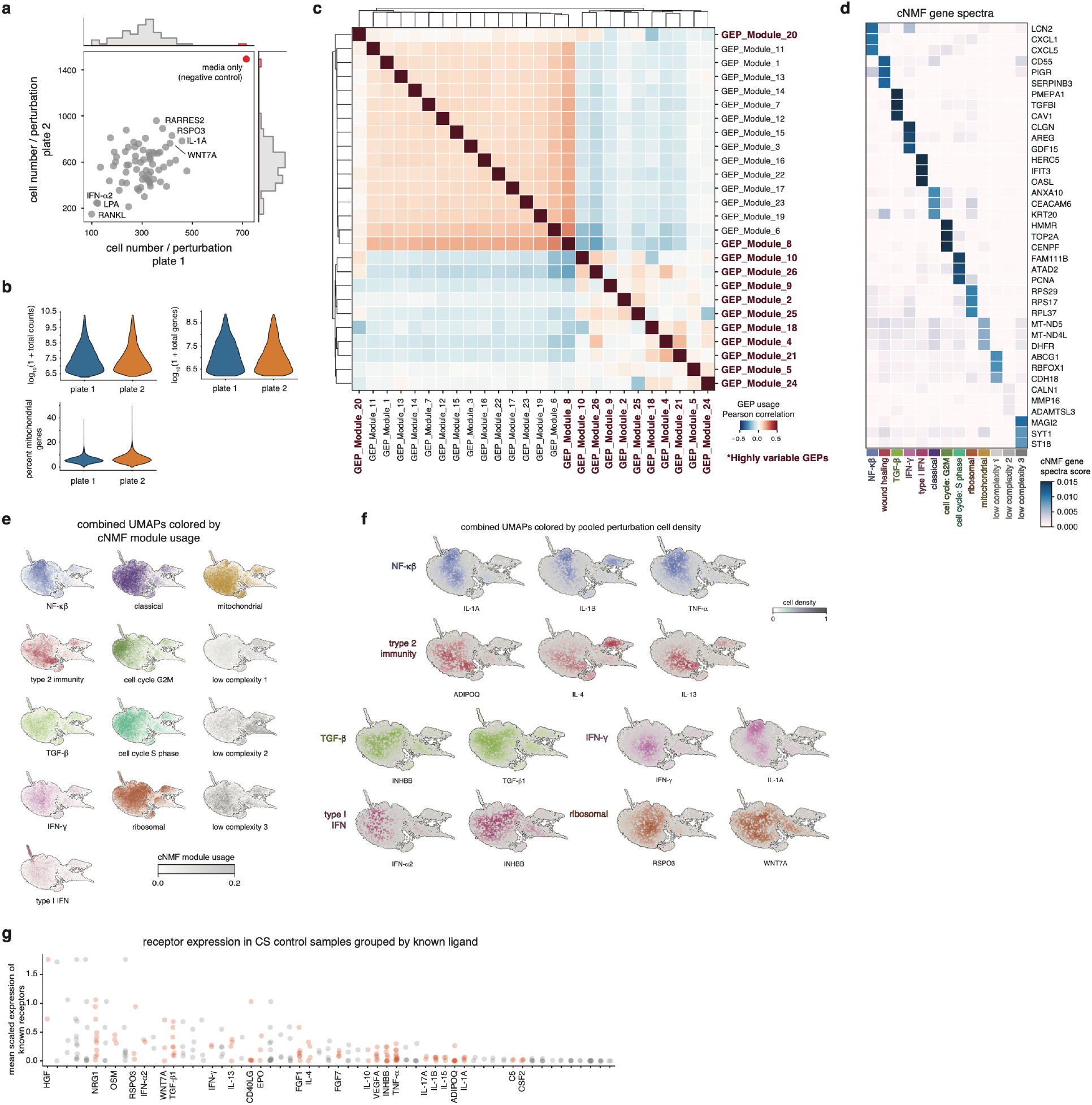
PDAC compressed screen scRNA-seq quality metrics and cNMF modules. **a**, Scatter plot of the number of cells per perturbation across all pools in each replicate plate. **b**, Violin plots of the number of UMIs, the number of unique genes, and the percent of genes that are mitochondrial in the compressed scRNA-seq dataset. **c**, Heatmap of the pairwise correlations of cNMF modules by usage across cells. **d**, Top three genes by gene spectra score for the highly variable cNMF modules. **e**, UMAP visualization all cells from both compressed screens, colored by cNMF module usage. **f**, UMAP visualizations all cells from both compressed screens, colored by density of cells from pools containing specific ligands. **g**, Ordered scatter plot of mean cognate receptor expression for each screened ligand over control PDAC cells in the compressed scRNA-seq dataset, colored by ligands with significant effects on identified cNMF GEPs.

**Extended Data Figure 3:**
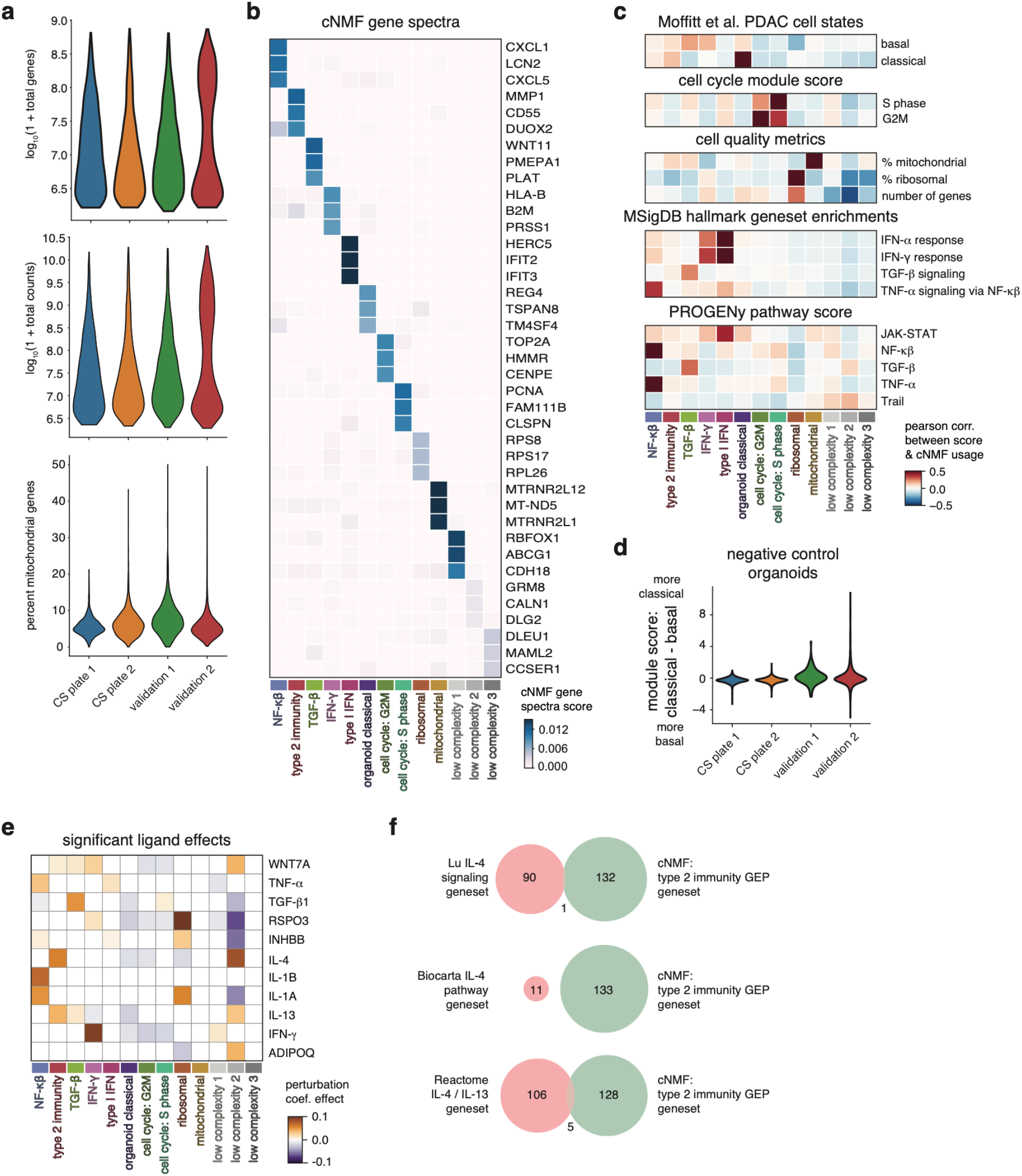
Single ligand perturbation experiment scRNA-seq quality metrics and cNMF modules. **a**, Violin plots of the number of UMIs, the number of unique genes, and the percent of genes that are mitochondrial in the single-ligand scRNA-seq dataset. **b**, Heatmap of the top three genes by gene spectra score for the single ligand cNMF modules that corresponded with the highly variable compressed cNMF modules. **c**, Heatmaps visualizing the Pearson correlation across cells of the usage of the select single-ligand cNMF gene expression programs and the module score for existing gene signatures. **d**, Violin plot of the Moffit classical module score – Moffit basal module score for all cells from organoids grown in media only from the different single ligand experiments. **e**, Heatmap of the non-zero regression coefficients by ligand for all single ligand cNMF modules corresponding with the highly variable cNMF modules from the compressed screen. **f**, Venn diagrams of the number of intersecting and unique genes between the cNMF type 2 immunity GEP and corresponding signatures in MsigDB.

